# Community assembly explains invasion differences between two contrasting forest types

**DOI:** 10.64898/2026.03.05.709929

**Authors:** Urmi Poddar, Tracey Dong, Kristi Lam, Vivianne Lee, Paul Wilson, Jessica Gurevitch, Rafael D’Andrea

**Affiliations:** Department of Ecology & Evolution, Life Sciences Building, Stony Brook University, Stony Brook, NY, USA – 11794; Roslyn High School, 475 Round Hill Rd, Roslyn Heights, NY, USA – 11577; Smithtown High School East, 10 School Street, St. James, NY, USA – 11780; Department of Forestry & Natural Resources, Purdue University, 715 W State St, West Lafayette, IN, USA – 47907

**Keywords:** Determinants of plant community diversity and structure, environmental filtering, functional traits, invasion ecology, pine barrens, temperate forests, species pools

## Abstract

Plant communities within a metacommunity can vary widely in their degree of invasion by introduced species. Disturbance, propagule pressure, and biotic resistance are common explanations for this variation, but empirical evidence for these hypotheses is mixed. Alternatively, the community assembly framework predicts that local assembly filters determine both native and exotic composition, but lower trait variation in the introduced species pool may exclude them from certain sites. We examined evidence for this framework using observational data from forests and woodlands of Long Island, NY, USA. These forests vary in vegetation composition and invasion along a soil gradient. They are also highly disturbed and fragmented, yet some stands have almost no introduced plants. Using data collected in 1998 and 2021-22, we quantified relationships between community composition, soil characteristics, and functional traits for native and exotic assemblages, as indicators of environmental filtering. We found similar trait-environment relationships in native and introduced species, suggesting that both groups follow the same local assembly rules. Introduced species were predominantly found in sites with more nutrient-rich soils and were absent from sites with nutrient-poor soils. At the regional scale, the exotic species pool was biased toward trait values favored in more nutrient-rich environments, particularly high growth rates and low leaf C:N ratios, which explains their absence from nutrient-poor environments. These patterns were consistent over time, and stands that were uninvaded in 1998 remained so in 2021-22, supporting the robustness and reliability of short-term studies. This study shows that invasion patterns in plant communities can be explained by the assembly rules that govern native species. By linking local environmental filtering with regional species pool characteristics, this work advances our understanding of how some communities remain uninvaded despite high disturbance and propagule pressure. Overall, these results highlight the utility of the community assembly framework, and emphasize the importance of regional processes in constraining the local distribution of introduced species.

## Introduction

Invasive alien species are a major threat to biodiversity and ecosystem functioning, and have negative economic impacts as well (Charles & Dukes, 2008; Gallardo et al., 2016; Mollot et al., 2017; Diagne et al., 2021). At the global scale, almost all parts of the world are threatened by introduced species (Early et al., 2016; Diagne et al., 2021). But at a smaller scale, some communities remain relatively uninvaded, even while others in the same metacommunity may be highly invaded. Gaining a better understanding of why certain communities are less invaded than others is important for identifying the drivers of invasion.

Several hypotheses have been proposed to explain the spatial distribution of introduced species at the regional or metacommunity scale. For example, the propagule pressure hypothesis states that communities receiving more exotic propagules should be more invaded (Lockwood et al., 2005; von Holle & Simberloff, 2005; Colautti et al., 2006; Cassey et al., 2018). Similarly, according to the disturbance hypothesis, disturbed communities are expected to be more invaded than undisturbed ones (Elton 1958; D’Antonio et al., 1999; Davis et al., 2000; Shea & Chesson, 2002; Mata et al., 2013; Jauni et al., 2015). Alternatively, biotic resistance by native communities may limit invasion (Levine et al., 2000; Mitchell et al., 2006), with more diverse communities predicted to be less invasible under the diversity–invasibility hypothesis (Elton, 1958; Levine, 2000; Beaury et al. 2020; Zhang, Liu, Brunel et al., 2020).

However, the patterns of exotic plant abundance and distribution across the forests of Long Island (Suffolk County), NY, USA appear to run counter to these hypotheses. A part of the New York City metropolitan area, Suffolk County has a high human population density (∼650 persons/km^2^ in 2020; U.S. Census Bureau, 2023) and a highly modified landscape dominated by low to medium density housing. Forest stands in this region are typically fragmented and surrounded by suburban development. They can be divided into two main community types: hardwood forests, and pine barrens (Conrad 1935; Greller, 1977; Jordan et al., 2003). Hardwood forests, found on more fertile, finer-textured soils, are characterized by closed canopies of mesic deciduous trees (Greller, 1977). The pine barrens, in contrast, occur in sandy, nutrient-poor soils (Kurczewski & Boyle, 2000; Jordan et al., 2003; Quigley et al., 2020), and are fire-tolerant communities with partially-open *Pinus rigida*-dominated canopies, and ericaceous shrub understories (Olsvig et al., 1998; Jordan et al., 2003). Despite their proximity to each other, these two forest types differ strikingly in their plant invasion levels (Fig. 1): the pine barrens are almost completely uninvaded, while hardwood forests range from moderately to heavily invaded (Howard et al., 2004).

**Figure 1:**
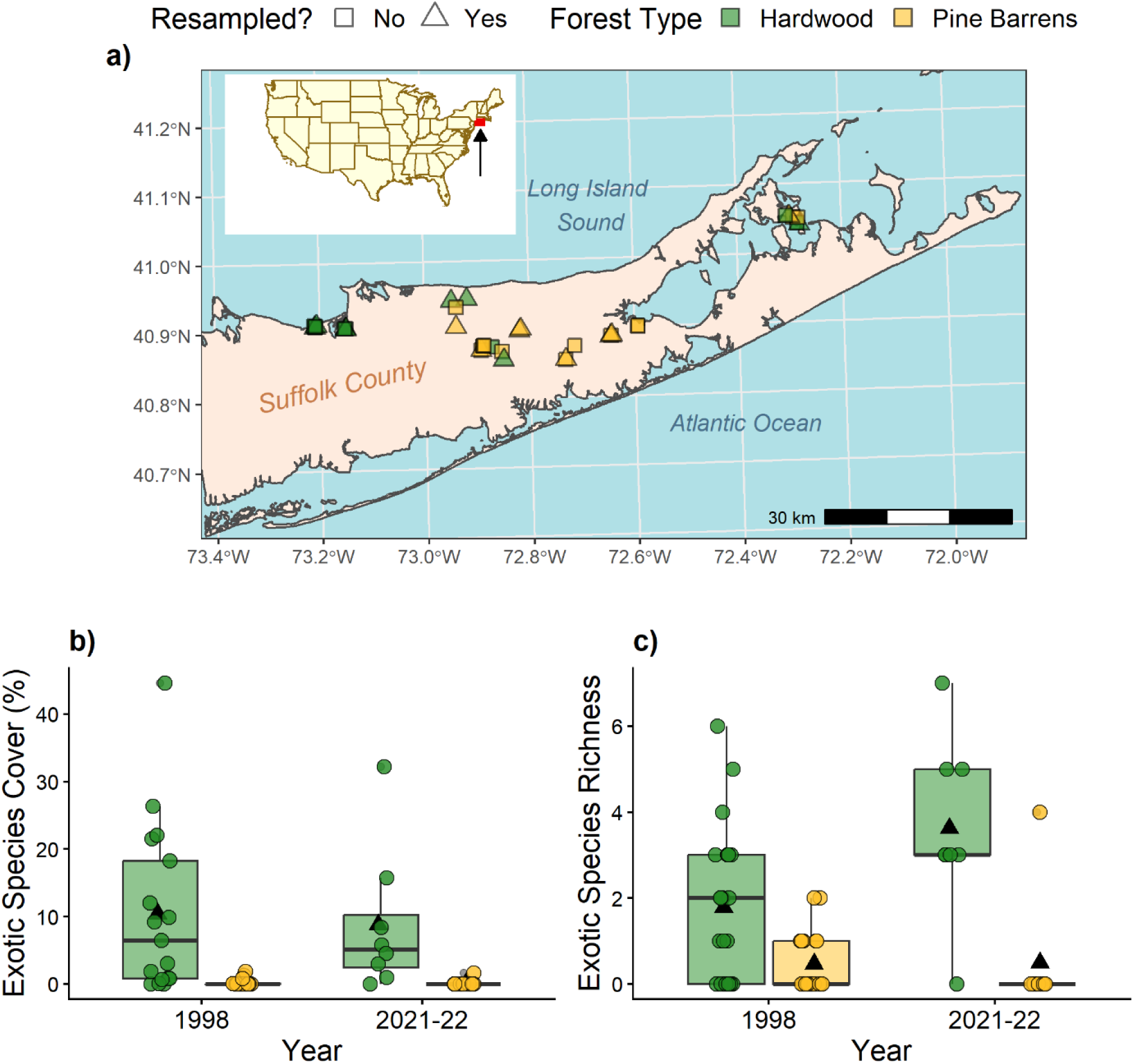
a) Map of Suffolk County, NY, USA, showing the location of study sites. Inset map shows the continental U.S. with a red bounding box (see arrow) around the study area. b-c) Exotic species cover (b) and richness (c) in the two forest types, in 1998 and 2021-22 (green - hardwood, yellow – pine barrens). Exotic species cover refers to the cover of all exotic plant species in a given site, divided by total cover of all species. Circles in b-c represent study sites, and black triangles denote the mean of each group.

This invasion pattern cannot be explained by propagule pressure or disturbance, as both forest types likely face similar levels of anthropogenic disturbance, fragmentation, and propagule pressure, and occur in close proximity, forming a spatial mosaic (Fig. 1a). Neither forest type is pristine; the high human density in the region results in substantial anthropogenic disturbance, which should make them both highly invasion-prone. The diversity-invasibility hypothesis also falls short, as hardwood forests have higher native plant diversity than the pine barrens (Howard et al., 2004), but are also more invaded. Thus, these patterns of invasion highlight the limits of current understanding of the drivers of invasibility. Moreover, the pine barrens are a globally rare and threatened ecosystem type (Kurczewski & Boyle, 2000; Sohl & Sohl 2012), and therefore, understanding their seemingly low invasibility may also aid in their conservation.

A previous study on this system (Howard et al., 2004) found that exotic plant cover was positively correlated with soil nutrient levels, both within and across the two forest types. This suggests that the nutrient-poor soils of the pine barrens may, in some way, contribute to their low degree of invasion. Similar patterns of low plant invasion in nutrient-poor soils or low-productivity environments have also been observed elsewhere (Huenneke et al., 1990; Alpert et al., 2000; Gerhardt & Collinge, 2007; Sardans et al., 2017), suggesting a link between ecosystem productivity and invasibility. However, the mechanisms preventing introduced plant establishment in low-productivity/low-nutrient environments remain unclear.

The community assembly framework for invasions (Shea & Chesson, 2002; Gallien and Carboni, 2017; Pearson et al., 2018; Brown & Barney, 2021) may provide an alternative and more general framework to explain these invasion patterns. According to this framework, introduced species are subject to the same local biotic and abiotic assembly filters that that determine native composition. Since these filters are agnostic to species provenance (Pearson et al., 2018), they should, at the local scale, influence both native and exotic composition in similar ways. At the regional scale, however, there may be differences in the trait pools of native and introduced species, due to introduction bias (van Kleunen et al., 2015; Kinlock et al., 2022) and differences in their evolutionary and migration history (Colautti et al., 2006; Fridley and Sax, 2014; Buckley and Catford, 2016). These regional differences constrain which species are available to pass through local assembly filters. Thus, according to the community assembly framework, the distribution of introduced species in a metacommunity should depend on the local assembly filters associated with each community, and the regional availability of introduced species that are adapted to these local filters. This provides a starting point for identifying invasion drivers using prior knowledge of local native assembly, which is particularly useful when conventional invasion hypotheses are insufficient, as seen in our system.

Although community assembly is driven by several processes (reviewed in Leibold et al., 2004; Vellend, 2016), here we focus on environmental filtering by the soil environment, along with regional pool differences, as this provides the most plausible hypothesis for the observed patterns of invasion in this study system. Specifically, Howard et al. (2004) found that both native composition and introduced plant abundance in these forests are correlated with soil characteristics, suggesting that the soil environment influences both native and introduced species composition. Therefore, we hypothesized that the introduced species distribution in our study system reflects soil-based environmental filtering, along with a lack of low nutrient-tolerance traits in the regional pool of introduced species (Fig. 2). We also hypothesized that the local composition of native and introduced species assemblages is determined by the same local environmental filters. Since environmental filters favor species with traits suited to prevailing conditions (Keddy, 1992; Liebold et al., 2004; Keddy & Laughlin, 2021), they are expected to give rise to measurable trait-environment associations (Münkemüller et al., 2012). Therefore, we looked for trait-soil associations as a signature of environmental filtering.

**Figure 2:**
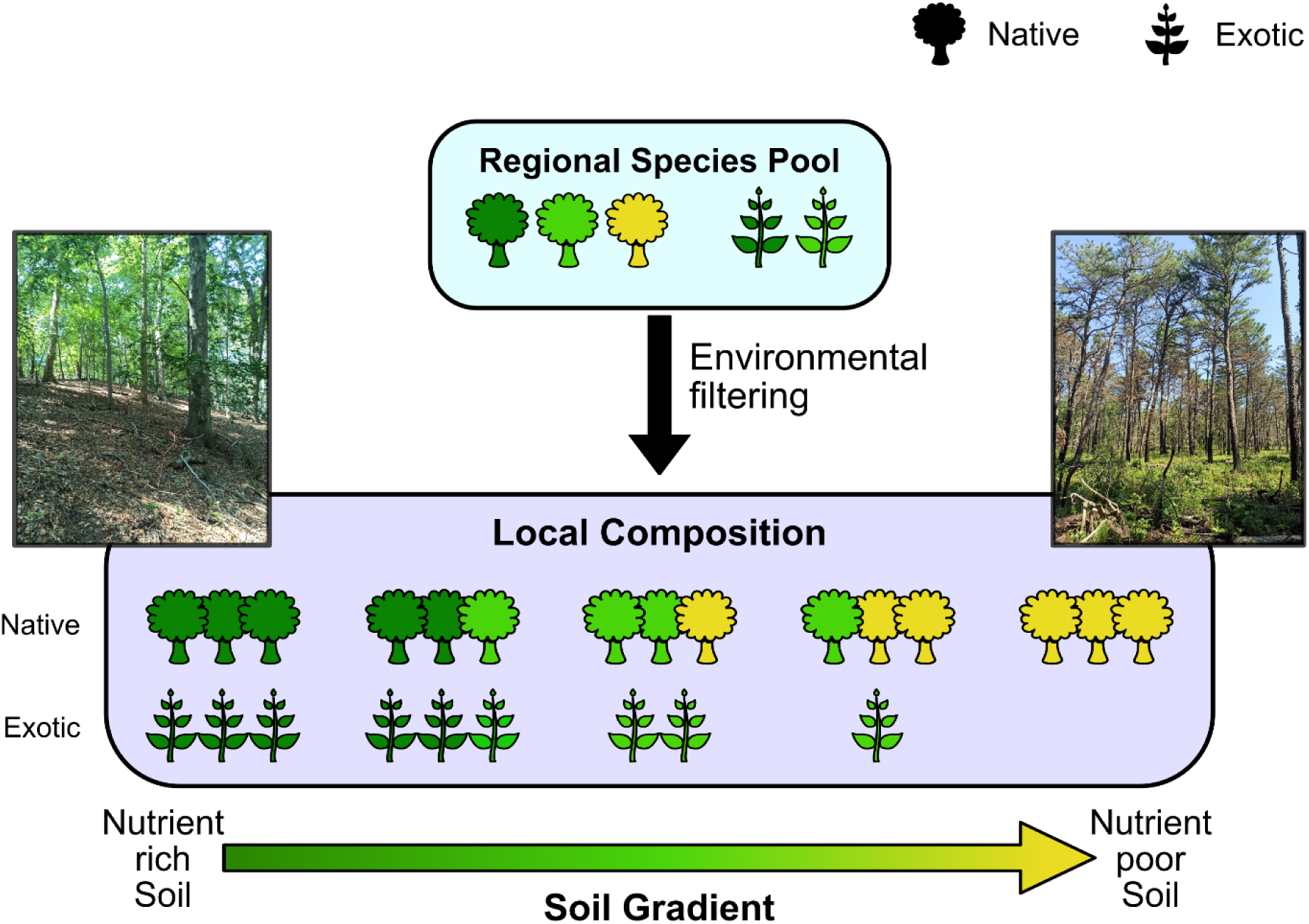
Hypothesized mechanism of native and exotic community assembly in Long Island’s forests. The environmental filtering model of community assembly predicts that local composition depends on the environment, with species being filtered from the regional species pool based on their functional traits. Here, the plant icons represent hypothetical species, with colors indicating different trait values, which determine their environmental (soil) preference. Taxonomic and functional composition of native and exotic species changes along the soil gradient as a result of environmental filtering by the soil. However, the regional species pool of exotic species lacks the trait values needed to survive in nutrient-poor soils, and therefore, their total abundance also changes along the soil gradient. Note that the shape of the native and exotic plant icons does not represent their life-form. Inset photographs show a representative hardwood forest (left), which are associated with more nutrient-rich soils, and a representative pine barrens forest (right), which are associated with nutrient-poor soils. Photo credit: The authors.

To identify the drivers of invasibility, the temporal dynamics of introduced species populations also need to be accounted for, as invasions often change in intensity and extent, typically expanding and increasing in dominance over time (Sakai et al., 2001; Pyšek & Hulme, 2005; Colautti & Barrett, 2013). Time lags in invasion, which are known to be common (Crooks & Soulé, 1999; Essl et al., 2012; Duncan, 2021) can further confound the study of invasibility. Communities that were previously uninvaded, and thus appeared to have low invasibility, may later become invaded (Essl et al., 2012). Moreover, native communities themselves are dynamic and may show temporal changes (Wu & Louks, 1995; Hubbel, 2001; Leibold et al., 2004; Franklin et al., 2016; Hu et al., 2022; Godoy et al., 2024), which could alter their invasibility. Therefore, we also asked whether the native and introduced plant composition of Long Island’s forests has changed over time.

In this study, we re-analyzed the vegetation composition and soil data from Howard et al. (2004; data collected in 1998) and carried out new vegetation surveys, in 2021 and 2022, for a subset of sites. We also gathered information on species functional traits using online databases. Using these data, first, we quantified soil-based environmental filtering by evaluating the relationship between community composition, functional traits and the soil environment across sites. We then tested whether native and introduced species show similar trait-soil relationships. Next, we looked for differences in the regional trait distributions of native and introduced species, and examined whether the introduced species pool lacked the trait values associated with nutrient-poor soils. Lastly, we looked for temporal changes in the taxonomic and functional composition of resampled sites, and in the degree of invasion of these sites, to assess whether the community assembly and invasion patterns of these forests were consistent across the 23 years between the studies.

## Methods

### Study sites

Community composition data used in this study was collected at two time points: 1998 and 2021-22. The 1998 data was gathered by Howard et al. (2004), who measured the vegetation composition and soil characteristics of 38 sites in Suffolk County, New York State, USA. The authors of the present study resampled 16 of these sites between 2021 and 2022.

Suffolk County is located on Long Island, east of New York City, and spans a land area of ∼2300 km^2^. It has a hot-summer humid continental climate (Dfa) (Beck et al., 2018). The soils of the study area are generally acidic and well-drained to excessively well-drained (USDA-SSA & CAES, 1975). All study sites were located in protected forest preserves, except for a few that were on the campus of Brookhaven National Laboratory. These stands are open to the public and consist of fragmented secondary forests with various land-use histories.

Pine barrens communities had partially-open canopies of *Pinus rigida* and *Quercus* tree species (mainly *Q. alba*, *Q. velutina,* and *Q. coccinea*), with dense understories dominated by shrubs such as *Q. ilicifolia*, *Vaccinium angustifolium*, *V. pallidum*, and *Gaylussacia baccata*. Hardwood forests, in contrast, were more diverse, with closed canopies dominated by *Acer rubrum*, *Betula lenta*, *Prunus serotina*, and *Quercus* species (mainly *Q. alba* and *Q. velutina*). Lianas such as *Toxicodendron radicans* and *Parthenocissus quinquefolia* were also common in hardwood forests. The understory tended to be sparse and varied from site to site in hardwood communities.

Details of the site selection process for the 1998 sites can be found in Howard et al. (2004). Sites in 2021-22 were selected to cover as much of the range of forest composition and invasion reported by Howard et al. (2004) as possible, as well as to maximize the distance between sites. Equal numbers of pine barrens and hardwood sites were chosen for resampling (for this purpose, forest type was determined by using the 1998 composition data and through visual assessments of potential sites).

### Vegetation composition

The methods used for quantifying vegetation composition were the same for both time points, detailed in Howard et al. (2004). In brief, a 30 m transect line was laid out in each site, and the cover of all vascular plants intersecting with the transect was noted. Ground cover (vegetation < 1 m height) was estimated through the line-intercept method, in 10-cm segments. Canopy cover was estimated visually at every 10-cm along the transect, using a modified periscope in 1998, and a GRS Densitometer (Geographic Resources Solutions, Arcata, California, USA) in 2021-22. Additionally, five 1 m x 1 m plots were placed within 5 meters of the transect line, and the percent cover of all vascular plant species rooted in each plot was noted. The World Flora Online (WFO) Plant List (https://wfoplantlist.org/; accessed June 2023) was used for taxonomic nomenclature. Species origin (native/introduced status) information was obtained from USDA Plants Database (https://plants.usda.gov/; USDA-NRCS, 2023).

The three cover estimates (canopy, transect-ground, plot) for each species were divided by their maximum possible values (100% cover) and then averaged to obtain a single abundance metric for each species in each site, which was used in all analyses. Prior to analysis, the abundance data for certain groups of species was combined due to difficulty in distinguishing between them in the field (these groups are henceforth referred to as species aggregates). This was done by summing the relative abundances of each individual species in an aggregate group, and then treating the aggregate as a single species. Species that were combined in this manner were: *Carya alba* and *Carya glabra*, *Elaeagnus angustifolia* and *Elaeagnus umbellata*, *Quercus rubra* and *Quercus velutina*, *Quercus coccinea* and *Quercus palustris*, *Solidago nemoralis* and unidentified *Solidago spp.*, *Vaccinium angustifolium* and *Vaccinium pallidum*, all *Vitis* species, and all unidentified graminoids. In most cases, the combined species were closely related and functionally similar, likely to hybridize in some cases (particularly *Quercus* species) and were often found together or in similar sites. Moreover, since our study is concerned with overall community-level patterns, we believe that this combining of a few species did not substantially impact our results.

Some of our analyses focused on comparisons of the two forest types (hardwood vs pine barrens). We used the following procedure to classify sites by forest type using vegetation composition. First, Ward’s hierarchical clustering, with Bray-Curtis dissimilarity, was applied to cluster sites based on similarities in composition. Then, the cluster dendrogram was cut to obtain two clusters. Finally, each cluster was assigned to a forest type based on the dominant native species of the sites in each cluster. After assignment of sites to forest type, the degree of association of each species with the pine barrens was calculated, as an indicator of species’ habitat preferences. For this, the cover of each species was summed across pine barrens sites and divided by its total cover across all sites. Note that a value of 1 indicates the given species was only found in the pine barrens, while 0 indicates that the species was only found in the hardwoods. Introduced species were expected to have values close to 0.

Using the R packages ‘vegan’ (Dixon, 2003), we ran an NMDS ordination, with Bray-Curtis dissimilarity, to visualize the variation in community composition across sites.

### Soil characteristics

Soil characteristics were measured by Howard et al. (2004) in 1998. In each site, they collected twelve soil cores from within a 10 m x 10 m square centered on the midpoint of the transect. These were air-dried and combined into a single sample per site, and then analyzed for pH, calcium, magnesium, cation exchange capacity (CEC), Bray-phosphorus, and soil texture (% sand, % silt, % clay). Further details can be found in the original study.

We standardized all soil variables by their means and variances prior to analysis. We carried out a PCA to visualize the variation in soil composition across sites, along with t-tests on each soil variable to test whether they significantly differed between the two forest types.

### Functional traits

Seven functional traits were used: expected height at maturity, specific leaf area (SLA), leaf carbon to nitrogen (C:N) ratio, shoot carbon to nitrogen (C:N) ratio (categorical), growth rate (categorical), fire tolerance (categorical), and life form (categorical) (Table 1). Height, specific leaf area (SLA), and growth rate are a part of the global plant economic spectrum (Diaz et al., 2015), i.e., the spectrum of plant life-history and competitive strategies. Leaf C:N ratio and shoot C:N ratio are measures of plant nutrient-use strategies (Ågren, 2004; Andersen et al., 2012; Reich 2014; Zhang, He, Liu et al., 2020), and they tend to vary along soil nutrient gradients (Luizão et al., 2004; Zhao et al., 2014; Gong et al., 2020; Peng et al., 2023). Fire tolerance measures the ability of a species to reestablish after a fire. It is expected to vary with forest type in this system, since pine barrens communities tend to be more fire-prone than hardwood communities. Life form refers to broad groupings of plant species based on similarities in physiognomy and life-history characteristics. Species belonging to the same life form are expected to respond similarly to the abiotic environment (Box 1996; Duckworth et al., 2000; Engemann et al., 2016).

**Table 1.** List of functional traits including in this study.

| **Trait** | **Categorical/**<br>**Continuous** | **Definition** | **Levels/range** |
| --- | --- | --- | --- |
| Fire tolerance | Categorical | Ability of the species to resprout, regrow, or reestablish from residual seeds after a fire. | None, low, medium, high |
| Growth rate | Categorical | Rate of increase in size of an individual after successful establishment. | Slow, medium, rapid |
| Life form | Categorical | Broad groupings of plant species based on similarities in physiognomy and life-history characteristics. | Herb, liana, shrub, tree |
| Shoot carbon: nitrogen (C:N) ratio | Categorical | Ratio of organic carbon to nitrogen contained in the above-ground biomass of each individual. | Low-medium, high |
| Height | Continuous | Expected height at maturity. | 0.2 – 57.9 m |
| Leaf carbon: nitrogen (C:N) ratio | Continuous | Ratio of organic carbon to nitrogen contained in each leaf. | 11.5 – 255.1 g/g |
| Specific leaf area (SLA) | Continuous | Leaf surface area per unit dry mass. | 1.6 – 53.4 mm<sup>2</sup>/mg |

Trait data was gathered from the TRY Plant Trait Database (https://www.try-db.org; Kattge et al., 2020) and the USDA Plants Database (https://plants.usda.gov/; USDA-NRCS, 2023). Height and leaf C:N ratio were log-transformed prior to analysis to reduce skew. Further details on the data source for each trait and trait data preprocessing can be found in Appendix S2.

Trait data was available for 60-90% of all species in this study. For each trait, species with available data represented 80-90% of total vegetation cover across all sites, except for leaf C:N ratio (∼70%) and life form (>99%). Trait interpolation was necessary for the dc-CA analysis (see next section), and was performed as detailed in Appendix S2.

### Relationship between community composition, functional traits, and soil environment

To evaluate the relationship between taxonomic composition, functional traits and the soil environment, we used double-constrained correspondence analysis (dc-CA). This is an ordination method that relates species composition to both functional traits and environmental predictors by maximizing the correlation between traits and environment (ter Braak et al., 2018). In practice, dc-CA first summarizes each site using canonical correspondence analysis (CCA), based on the traits of the species present, producing site scores that represent functional composition. These site scores are then related to environmental variables using redundancy analysis (RDA), to identify which environmental predictors that best explain variation in functional composition across sites. The resulting site scores, representing each site’s environment, are used to infer species’ environmental associations and the functional traits that best explain the same. The R package ‘douconca’ (ter Braak & van Rossum, 2025), was used for performing dc-CA, with separate analyses for 1998 and 2021-22. Model significance was assessed using the maximum permutation test (max test), which involves two independent permutation tests, one permuting species attributes (traits), and the other permuting site attributes (environmental variables). The more conservative (i.e. larger) p-value is then used for significance testing. This approach accounts for inflated type I error rates that often arise in trait-environment analyses (ter Braak et al., 2012; Zelený, 2018).

Due to multicollinearity between traits, we selected a reduced subset of traits for each time point, by performing CCA on the transposed composition matrix followed by stepwise selection, using the R package ‘vegan’ (Dixon, 2003). Lifeform was one of the traits selected through this procedure, but some of its levels showed high variance inflation factor (VIF) values in the dc-CA. Therefore, it was simplified to a binary liana vs. non-liana trait. All soil predictors except % silt and % clay were included. The latter two were excluded as the three soil texture variables (% silt, % clay and % sand) are strongly correlated with each other and sum to 100%; hence % sand was the only soil texture variable used.

To assess the similarity in the trait-environment relationships of native and introduced species, first, we visually compared the dc-CA scores of the two groups. The scores of introduced species were expected to be similar to those of hardwood-associated native species but dissimilar from pine barrens-associated native species, which would indicate that exotic species’ traits and environmental preferences align with those of co-occurring native species. Note that dc-CA produces two sets of species scores, one representing species traits, and the other representing their environmental preferences. We also tested whether species origin (native/introduced status) explained species’ environmental preferences beyond what can be explained by functional traits. For this, a second set of dc-CA models was fitted with species origin as a predictor and functional traits as conditioning variables. This allows for testing the effect of species origin after controlling for potential trait differences between native and introduced species. Therefore, if such a model is statistically significant, it would suggest that the environmental associations of introduced species are dissimilar to those of functionally-similar native species.

### Species pool differences

To compare the regional species pool trait distributions of native and introduced species, we first defined the regional pool as all species observed in either survey (1998 or 2021-22). We then constructed species pool trait distributions by assuming equal abundance of all species in the regional pool. Separate distributions were created for native and introduced species, which were then compared using two-sample Kolmogorov Smirnov tests (continuous traits) and Chi-squared tests of independence (categorical tests). For continuous traits, t-tests and F-tests were also used to compare means and variances, respectively.

### Temporal trends in composition

To test whether the taxonomic composition of the study sites significantly changed over time, we carried out PERMANOVAs (permutational MANOVA) on the composition of resampled sites, with sampling year (1998 vs 2021-22) as the predictor. Permutations were restricted to within site, i.e. the year labels were permuted within each site, to account for non-independence between samples from the same site. Two sets of PERMANOVAs were carried out, one on taxonomic composition, and another on functional composition. Additionally, we first carried out this analysis using all species, and then separately for native and introduced species to allow comparison between the two. Thus, there were six PERMANOVAs in total. Bray-Curtis dissimilarity was used for the PERMANOVAs on taxonomic composition. For the functional composition PERMANOVAs, the response variable was calculated as follows: first the functional distance between all pairs of species in the dataset was calculated, using the Gower distance metric, through the R package ‘cluster’ (Maechler et al., 2023), with all seven traits. Then, mean pairwise functional dissimilarity between sites, and between the two samples from each site, was calculated using the R package ‘picante’ (Kembel et al., 2010). This was then used as the response variable in the PERMANOVA.

To test whether the degree of invasion in the two forest types significantly changed over time, two mixed-effects ANOVAs were performed, one on log-transformed exotic species richness and another on square root-transformed introduced species cover per site. Forest type and sampling year were the fixed-effects predictors, and site was the random effect. The R package ‘lme4’ (Bates et al., 2003) was used for fitting the mixed-effects model.

All analyses were performed in R version 4.2.0 (R Core Team, 2022).

## Results

### Study sites

Taxonomic composition of the study sites could be divided into two distinct clusters, corresponding to the two forest types (Fig. S1). The cover and richness of introduced species was negligible in the pine barrens, while hardwood sites had significantly higher invasion; a pattern that persisted over time (Fig. 1, Table S1). Native species richness was higher in the hardwood sites than the pine barrens (Fig. S2), while the pine barrens tended to have higher ground cover and lower canopy cover than hardwood sites (Fig. S3).

Soil composition varied both within and across forest types. In the PCA of all soil variables, the first principal component axis was most strongly correlated with % sand, and the second axis was most strongly correlated with magnesium and calcium (Fig. S4). Pine barrens communities occurred across the range of soil textures, while hardwood forests were mainly found on moderately sandy soils. The two forest types differed mostly along the second axis, corresponding to differences in soil calcium and magnesium concentrations. Pine barrens mostly occurred on soils with the lowest Ca and Mg, while hardwood forests were not found on sites with higher Ca and Mg. This was further confirmed through the t-tests on individual soil variables, which showed that the two forest types differed significantly in soil calcium and magnesium, but not in any other soil variable (Table S2).

### Relationship between community composition, functional traits, and soil environment

In the dc-CA analysis, in both years, native species tended to segregate based on their forest type preferences, forming a pine barrens-to-hardwood gradient along the first two dc-CA axes (Figs. 3-4). Introduced species, particularly the most abundant ones, tended to be found at the hardwood-end of this gradient, suggesting that their traits and environmental relationships were similar to hardwood-specialized native species. These patterns were particularly distinct in species’ environmental preferences (Figs. 3a, 4a), while traits produced a less clear, though still noticeable, forest type-association gradient (Figs. 3b, 4b). Note that species’ forest type preferences were not used as predictors in the dc-CA models; rather these patterns emerged naturally from the functional traits and soil predictors included in the model.

**Figure 3:**
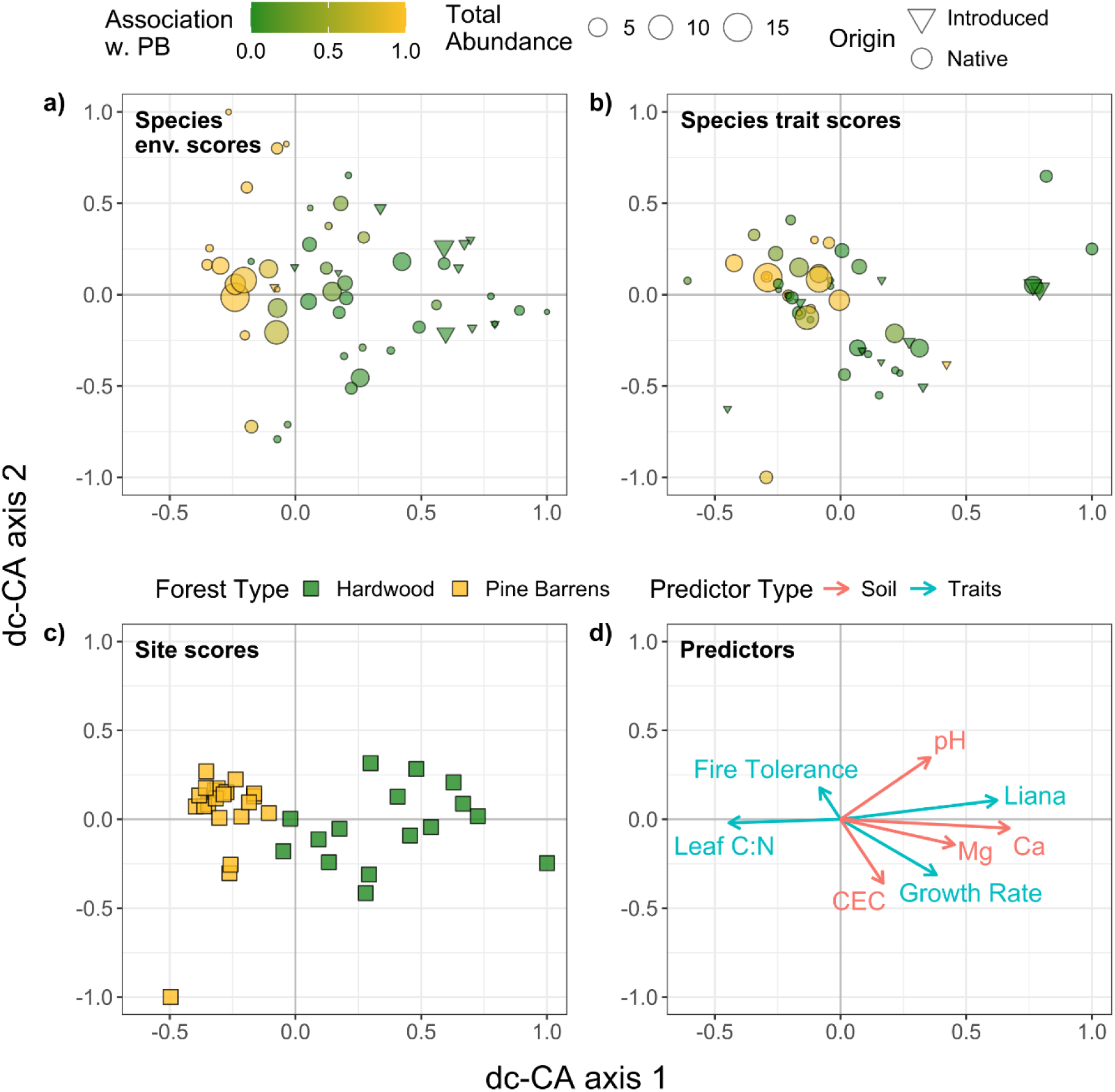
Ordination diagrams from the dc-CA on the 1998 dataset. a) Species scores representing species’ environmental preferences, b) species scores representing their functional traits, c) site scores representing functional composition, d) functional traits and environmental predictors. Colors in a) and b) denote the degree of association of each species with the pine barrens (0 – exclusively found in hardwoods, 1 – exclusively found in pine barrens), size of points indicates each species’ total abundance (sum of cover across sites), and shape indicates species origin. Only the most important traits and soil predictors are shown in d). CEC – cation exchange capacity.

**Figure 4:**
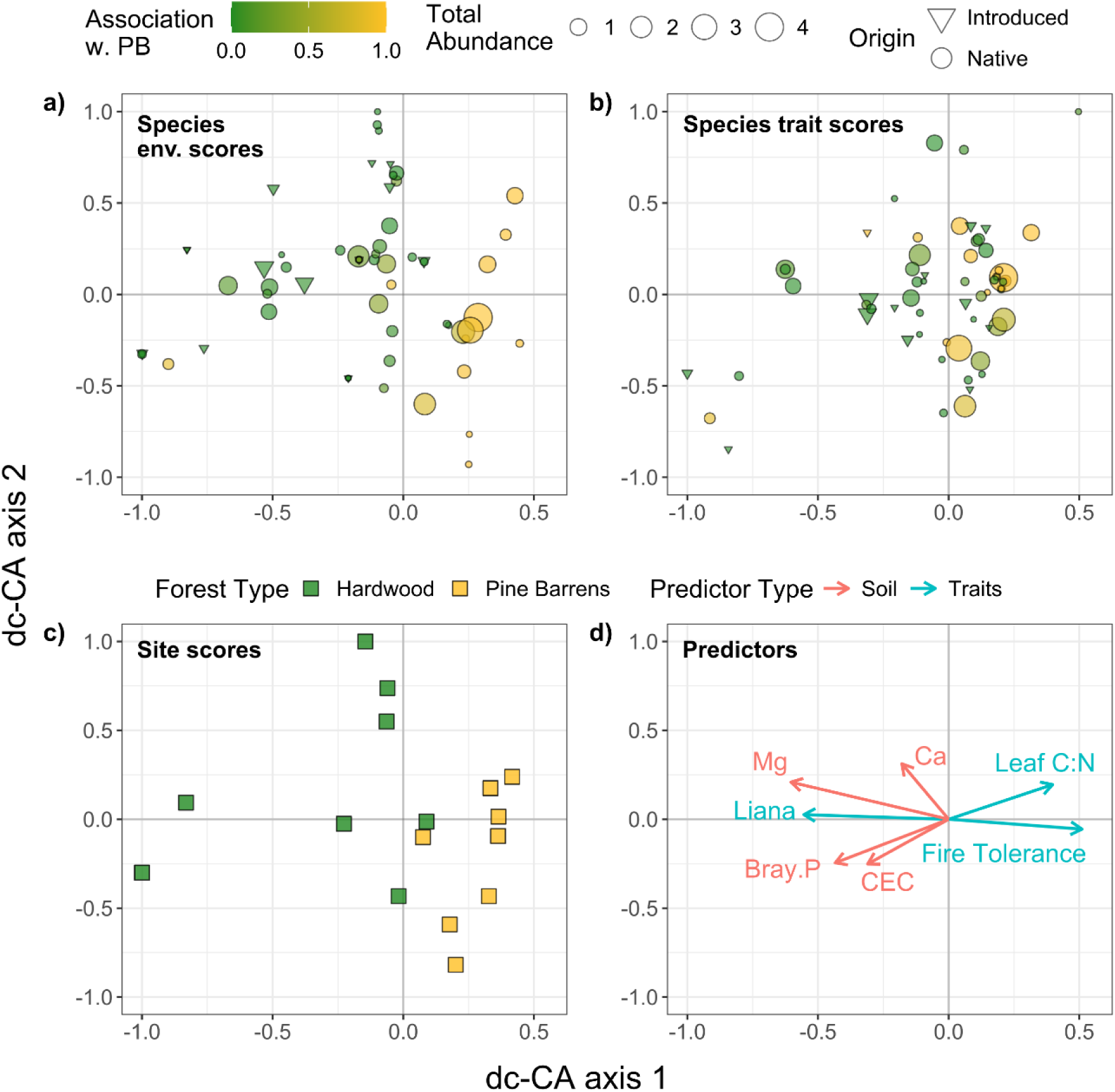
Ordination diagrams from the dc-CA on the 2021-22 dataset. a) Species scores representing species’ environmental preferences, b) species scores representing their functional traits, c) site scores, representing functional composition, d) functional traits and environmental predictors. Colors in a) and b) denote the degree of association of each species with the pine barrens (0 – exclusively found in hardwoods, 1 – exclusively found in pine barrens), size of points indicates each species’ total abundance (sum of cover across sites), and shape indicates species origin. In d) only the most important traits and soil predictors are shown. Bray.P – Bray phosphorous, CEC – cation exchange capacity.

In 1998, leaf C:N ratio, fire tolerance, growth rate and the liana trait were the traits selected for inclusion. Functional traits alone explained 29% of the variation in species distributions, while soil predictors alone explained 22% of the variation in species composition across sites. Traits and soil predictors combined explained 9% of the variation in community composition. The first dc-CA axis was statistically significant (Table S4), describing a gradient from low to high values of calcium and magnesium, along with decreasing leaf C:N ratio and increasing growth rate and prevalence of lianas. Pine barrens sites and pine barrens-associated species tended to have low scores on axis 1, while the opposite was true for hardwood sites and hardwood-associated species, including introduced species. The overall dc-CA model was statistically significant (Table S3).

In 2021-22, functional traits and soil predictors explained, respectively, 36% and 48% of the variation in community composition, out of which 20% was explained by traits and soil combined. Leaf C:N ratio, fire tolerance, SLA and liana were the traits included in the model. The first dc-CA axis was statistically significant (Table S4), and it was mainly correlated with decreasing soil magnesium and liana prevalence, and increasing fire tolerance and leaf C:N ratio. Hardwood sites and hardwoods-associated native species, as well as introduced species, tended to have low axis 1 scores, while the opposite was true for pine barrens sites and pine barrens-associated species. The overall dc-CA model was statistically significant (Table S3).

Models with species origin as a predictor and traits as conditioning variables were not statistically significant (Table S3), implying that native and introduced species did not differ in their trait-environment relationships.

### Species pool differences

The species pool trait distributions of native and introduced species significantly differed from each other for leaf C:N ratio and height (Tables S5-S8). In the case of height, the native species pool showed a bimodal distribution, with intermediate-height species being rare, while the introduced species pool showed a unimodal distribution centered on intermediate height values (Fig. 5a). The introduced species pool also had a significantly lower mean leaf C:N ratio than native species (based on t-test; Table S6, Fig. 5b). Additionally, the introduced species pool had a greater prevalence of rapid and medium growth rate species, while the native pool had a higher proportion of slow-growing species (Fig 5e), although this difference was not statistically significant.

**Figure 5:**
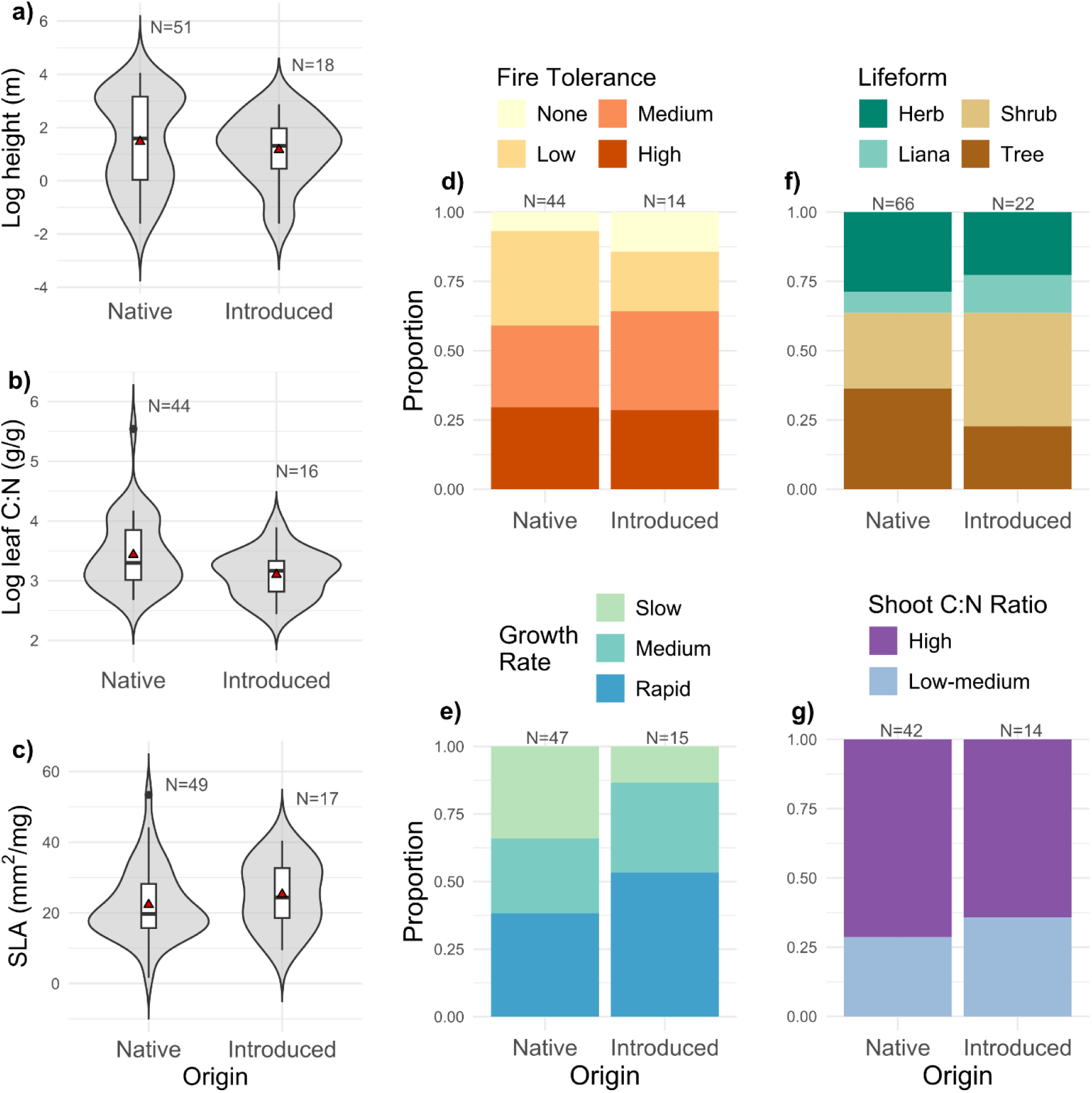
Species pool trait distributions of native and introduced species. a) log height, b) log leaf C:N ratio, c) specific leaf area (SLA), d) fire tolerance, e) growth rate, f) life form g) shoot C:N ratio. Red triangles inside the boxplots in panels a-c depict the mean values of the respective distributions.

### Temporal trends in composition

The taxonomic composition of resampled sites significantly changed over time, but functional composition did not (Table 2). This was seen in both native and introduced species assemblages (Table S9). The direction of change taxonomic composition was idiosyncratic (Fig. S1), suggesting no specific species consistently declined or increased across sites. Examinations of the changes in the cover of individual species further confirmed that the direction of change varied across species and sites (Fig S5, S6). Similarly, the cover and richness of exotic species showed no significant temporal changes (Fig. 1, Table S1). However, overall canopy cover and ground cover decreased over time in both forest types, with particularly large decreases in the latter (Fig. S3)

**Table 2.** Results of permutational MANOVAs testing whether the taxonomic and functional composition of resampled sites significantly changed over time.

| Taxonomic composition |  |  |  |  |
| --- | --- | --- | --- | --- |
| Predictor | Df | R<sup>2</sup> | F | P-value |
| Sampling year | 1 | 0.05 | 1.63 | **0.001\*** |
| Residual | 30 | 0.95 |  |  |
| Functional composition |  |  |  |  |
| Predictor | Df | R<sup>2</sup> | F | P-value |
| Sampling year | 1 | 0.03 | 0.87 | 0.24 |
| Residual | 30 | 0.97 |  |  |

\*p < 0.05

## Discussion

We found that the pine barrens of Long Island, a rare and threatened ecosystem type (Kurczewski & Boyle, 2000; Sohl & Sohl 2012), have remained largely uninvaded by introduced plants, despite continued anthropogenic disturbance and proximity to established invasive species populations. To understand the low invasion of these forests, we compared them to nearby hardwood forests which show varying degrees of invasion. Across both forest types, vegetation composition and invasion varied along gradients of soil nutrients, particularly magnesium and calcium. Slow-growing, fire-tolerant species with higher leaf C:N ratios tended to be found in the nutrient-poor soils of the pine barrens, while faster growing species with lower fire tolerance and leaf C:N ratios were associated with the higher nutrient soils of the hardwoods (Figs. 3-4). These trait-soil relationships did not significantly differ between native and introduced species (Table S3). Rather, the regional species pool of introduced species was biased towards trait values associated with the hardwoods, particularly low leaf C:N ratio and faster growth rates (Fig. 5), which likely explains their absence in the pine barrens. These patterns of community assembly and invasion were similar between the two time points, even though they were separated by more than two decades.

The trait-environment relationships identified here are consistent with previous research on plant life-history strategies and response to abiotic filters. In particular, we found a turnover from species with resource-conservative traits, such as slow growth rate and high leaf C:N ratio, to those with resource-acquisitive traits, such as high growth rate and low leaf C:N ratio, with increasing soil nutrients. This likely represents a trade-off between maximum potential growth rate and nutrient-use efficiency (Lambers & Poorter, 1992; Reich 2014), and is consistent with other studies on plant trait responses to soil nutrients (Ågren, 2004; Andersen et al., 2012; Reich 2014; Zhang, He, Liu et al., 2020; Peng et al., 2023). Similarly, the observed association of lianas with nutrient-rich soils can be explained by their tendency to possess resource-acquisitive life history strategies (Gallagher & Leishmanm, 2012; Collins et al., 2016). Additionally, the negative relationship between fire tolerance and soil nutrients is likely due to the fire-prone nature of the pine barrens, which results in more fire-tolerant species being found in nutrient-poor soils. Interestingly, we also observed greater trait variation among hardwood-associated species than pine barrens-associated ones. This suggests that species with varying ecological strategies can coexist in the hardwoods, while a narrower set of ecological strategies are selected for in the pine barrens. Whether this is driven by the soil environment or by other assembly processes is not known.

These trait-environment relationships did not significantly differ between native and introduced species, indicating that the two groups follow the same assembly rules at the local scale. Introduced species showed a preference for high soil calcium and magnesium, similar to hardwoods-associated native species, and they also tended to possess traits similar to the latter. Given that introduced species were almost exclusively found in the hardwoods, their traits and environmental preferences were thus similar to those of co-occurring native species. A few other studies have also identified similarities in the assembly of co-occurring native and introduced species (Meiners et al., 2009; Lemoine et al., 2015; Pearson et al., 2018; Poddar et al., 2024), supporting the idea that all species in a community are subject to the same local assembly filters. Interestingly, this association of introduced species with the more-diverse hardwood communities contradicts the well-known biotic resistance hypothesis (Elton, 1958; Levine, 2000; Catford et al., 2019; Beaury et al. 2020; Su et al., 2023). Instead, these results are consistent with the lesser-known biotic acceptance hypothesis (Stohlgren et al., 2006; Souza et al., 2011), which posits that environments that are favorable to a large number of native species, and thus have high native diversity, should also be favorable to introduced species, resulting in positive diversity-invasion relationships.

These habitat preferences of introduced species can be further explained by the species pool comparison. This comparison showed that at showed that at the regional scale, introduced species tended to possess resource-acquisitive traits such as high growth rate and low leaf C:N. This likely limited the range of habitats they could occupy locally. Although most conventional hypotheses on community invasibility, such as the diversity-invasibility hypothesis (Elton, 1958), the disturbance hypothesis (Elton, 1958), and the fluctuating resources hypothesis (Davis et al., 2000), focus on local assembly filters, this study, along with a few others (Meiners et al., 2009; Duffin et al., 2019; Pearson et al., 2023; Poddar et al., 2024), demonstrates the importance of regional processes in explaining local invasion patterns. Regional filters associated with the migration and introduction history of introduced species constrain the availability of species adapted to a given local ecosystem, thus influencing the local distribution and composition of introduced species. From a conservation perspective, preventing the invasion of currently-uninvaded ecosystems, particularly those associated with stressful environments, may thus require regional-scale efforts to restrict the introduction of species adapted to those environments.

These patterns of invasion and community assembly were remarkably consistent over a 23-year period. Many previous studies on community assembly (e.g. Kraft et al., 2008; Mori et al., 2013; da Silva et al., 2018; Fichaux et al. 2019; Jiao et al., 2020) and invasion (e.g. Funk & Vitousek, 2007; Andersen et al., 2012; Su et al., 2023; Westerband et al., 2025) have relied on data from a single time point. Whether the results of such studies would hold across time is not known, given the dynamic nature of plant communities (Wu & Louks, 1995; Hubbel, 2001; Leibold et al., 2004; Franklin et al., 2016; Hu et al., 2022; Godoy et al., 2024) and of invasive species populations (Crooks & Soulé, 1999; Essl et al., 2012; Duncan, 2021). Here we found that the assembly processes in Long Island’s forests remained largely similar after 2 decades. Similarly, the degree of invasion of these sites did not change significantly, and the low degree of invasion in the pine barrens reported in 1998 was also seen in the later data. Even in the invaded hardwood forests, there were no significant increases in the degree of invasion despite the presence of established invasive species in 1998 and continued high levels of fragmentation and disturbance. Although the taxonomic composition of these forests did significantly change over time, functional composition did not, indicating that the former represents functionally-neutral fluctuations in species composition. This lack of change is reassuring, as gathering long-term data can be challenging.

But in sharp contrast to community composition, both total ground cover and total canopy cover declined over time (Fig. S3). Ground cover in particular showed large decreases. This is concerning, as it indicates a decline in average plant population sizes, which may affect ecosystem functioning and threaten rare and endangered plant species. Such declines could be caused by the high overpopulation of white-tailed deer (*Odocoileus virginianus*), an overabundant herbivore that has hampered plant recruitment in many North American forests (Rooney & Waller, 2003; McShea, 2012; Russell et al., 2017). However, the exact causes and consequences of this decline cannot be determined in the present study.

One potential limitation of this study is that the regional pool trait distributions were constructed from the traits of observed species. This method likely underestimates the variation in the regional pool, as our field sampling methodology is not meant to provide comprehensive species lists of the studied sites (Howard et al., 2004). However, as our species pool analysis focused on relative differences between native and introduced species, rather than on absolute trait variance, and our sampling methodology is not expected to be biased towards either group of species, we believe our conclusions are robust to this limitation. Additionally, this study does not consider assembly filters other than soil-based filtering and species pool effects. In particular, biotic interactions such as competition are also known to be important drivers of invasibility (Levine et al., 2000; Mitchell et al., 2006; Beaury et al. 2020; Su et al., 2023). These interactions could not be included in the present study due to the difficulty of quantifying biotic interactions with observational data. Moreover, the pine barrens have shorter fire return intervals than the hardwood forests (Kurczewski & Boyle, 2000; Jordan et al., 2003), and fires tend to remove pre-existing vegetation, which should reduce the competition faced by potential invaders. The fact that the pine barrens are largely uninvaded despite their fire-prone nature suggests that competition may play a lesser role in their invasion resistance. Nevertheless, the role of competition and other biotic interactions cannot be completely ruled out, and their contributions to the observed patterns of invasion need to be further studied.

