## Supplement for "Community assembly explains invasion differences between two contrasting forest types"

**Appendix S1: Additional Figures & Tables**

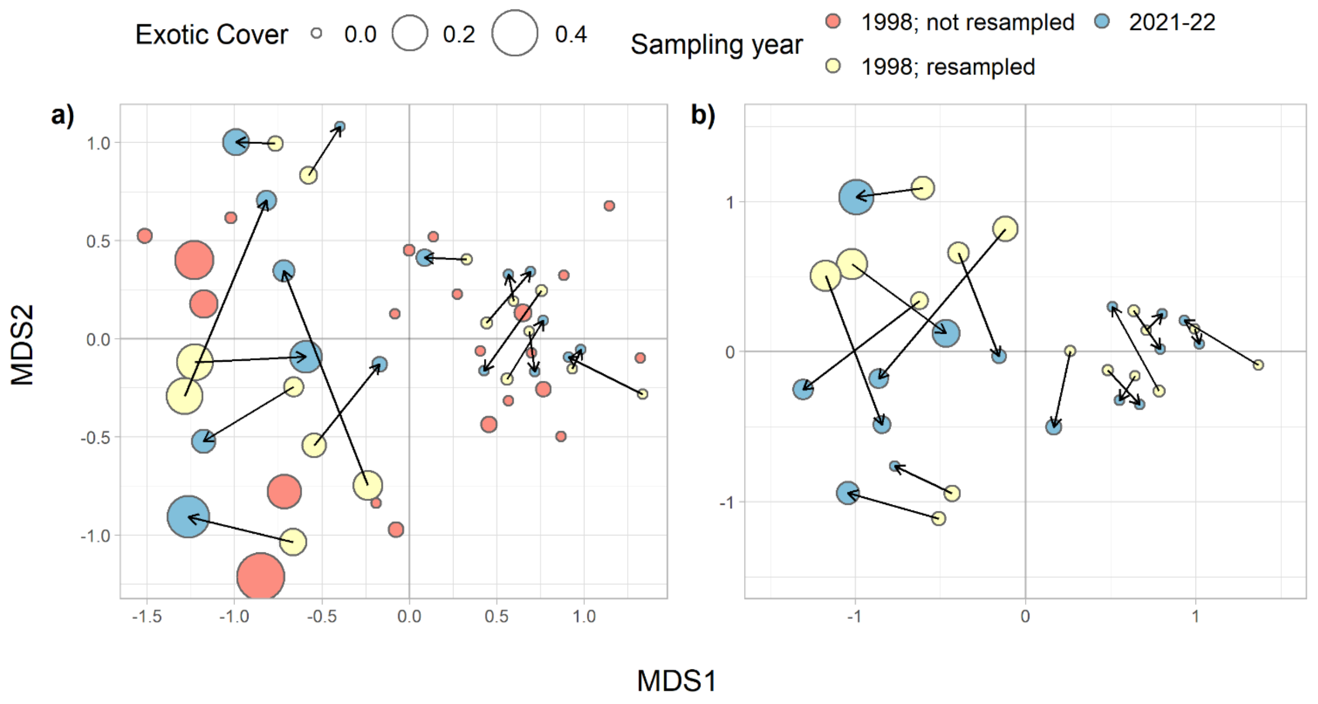

**Figure S1:** NMDS of vegetation composition of a) all sites and b) resampled sites only, showing two visually distinguishable clusters of sites in both cases. Sites with low axis 1 scores correspond to hardwood forests and those with high scores correspond to the pine barrens. (Note that sites were classified by forest type through Ward clustering, not through visual examinations of NMDS scores. However, separation of sites along NMDS axis 1 aligned well with the clustering-based classification.) The size of the points depicts the relative cover of introduced species in each site as a proportion of total vegetation cover. Arrows link 1998 data to the corresponding 2021-22 data from the same location, in order to depict change in composition over time. The idiosyncratic direction of the arrows indicates no specific species increased or decreased across all times.

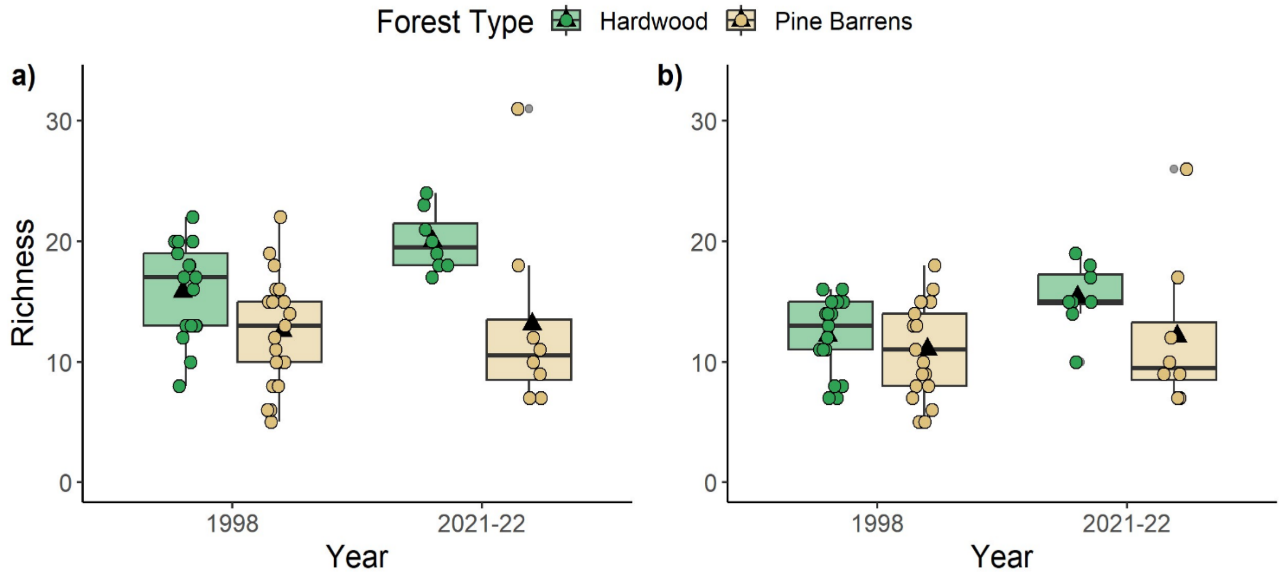

**Figure S2:** Species richness of a) all species and b) native species in the two forest types, across time.

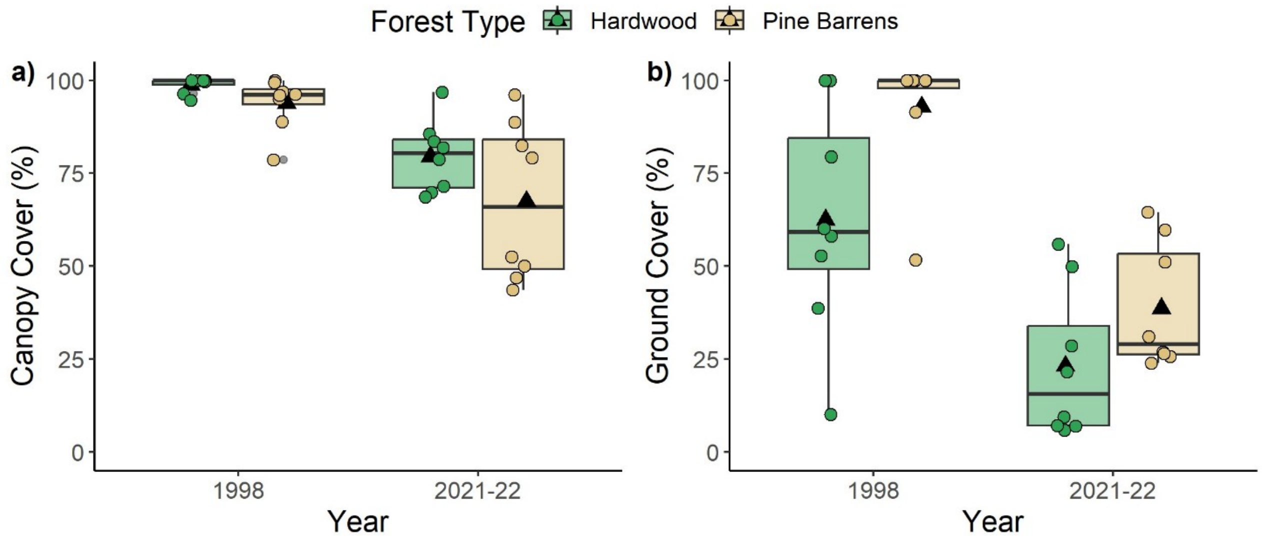

**Figure S3:** Percentage canopy cover (a) and ground cover (b) in the two forest types, across time. From the 1998 dataset, only sites that were resampled are shown.

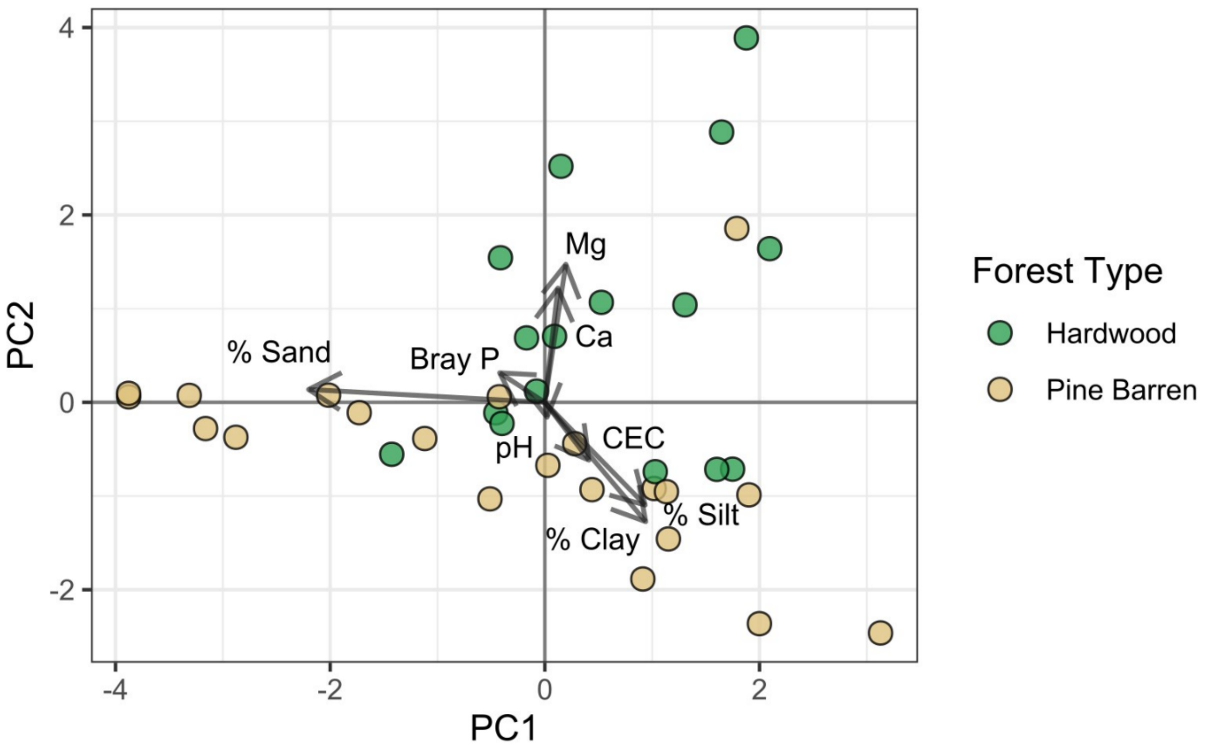

**Figure S4:** PCA of soil variables, showing differences within and across forest types. Bray P = Bray phosphorous, Ca = calcium, CEC = cation exchange capacity, Mg = magnesium.

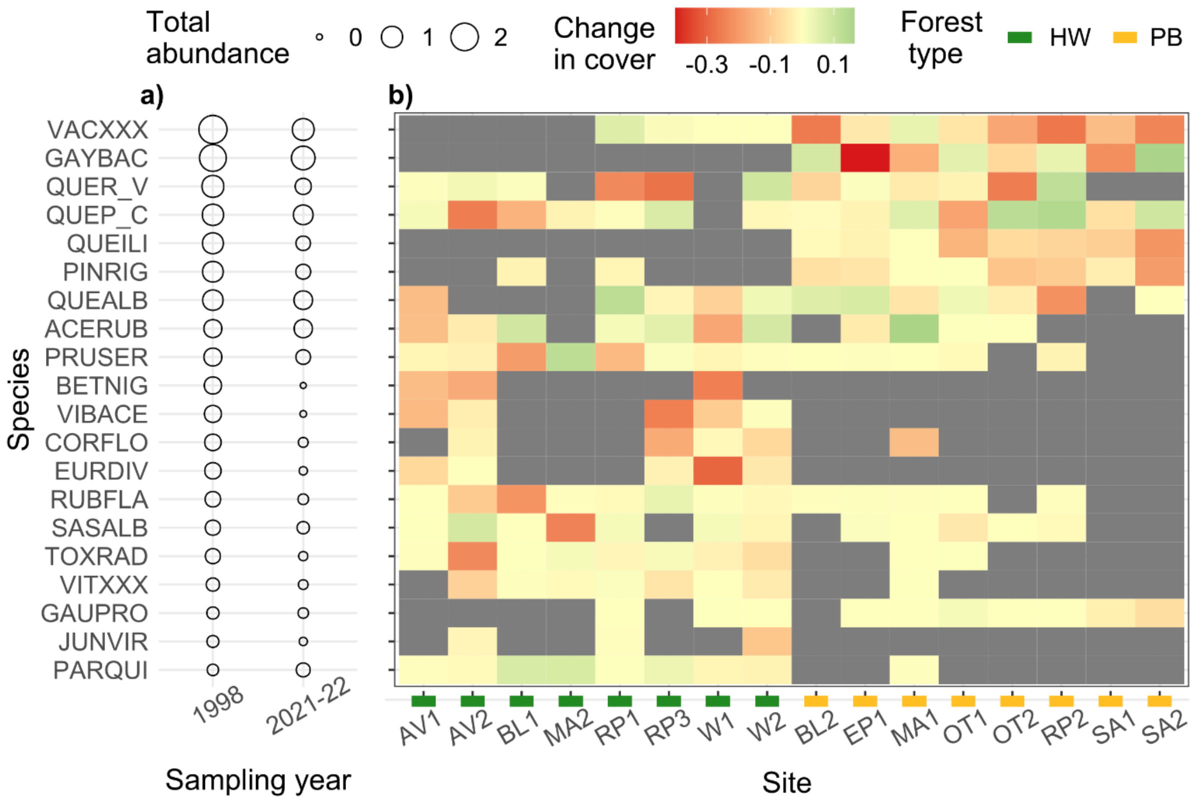

**Figure S5:** Temporal changes in the abundance of the 20 most common native species in 1998. a) Total abundance (sum of cover across sites) of each species in 1998 and 2021-22. b) Change in cover of each species in each site (cover in 1998 – cover in 2021-22). Sites are arranged by forest type. Positive values (light green) indicate increase, and negative values (red) indicate decrease over time. Grey fill indicates absence of the given species in the given site at both time points. From the 1998 dataset, only the data from resampled sites was used. Species names associated with the six-letter codes in a) are shown in table S10.

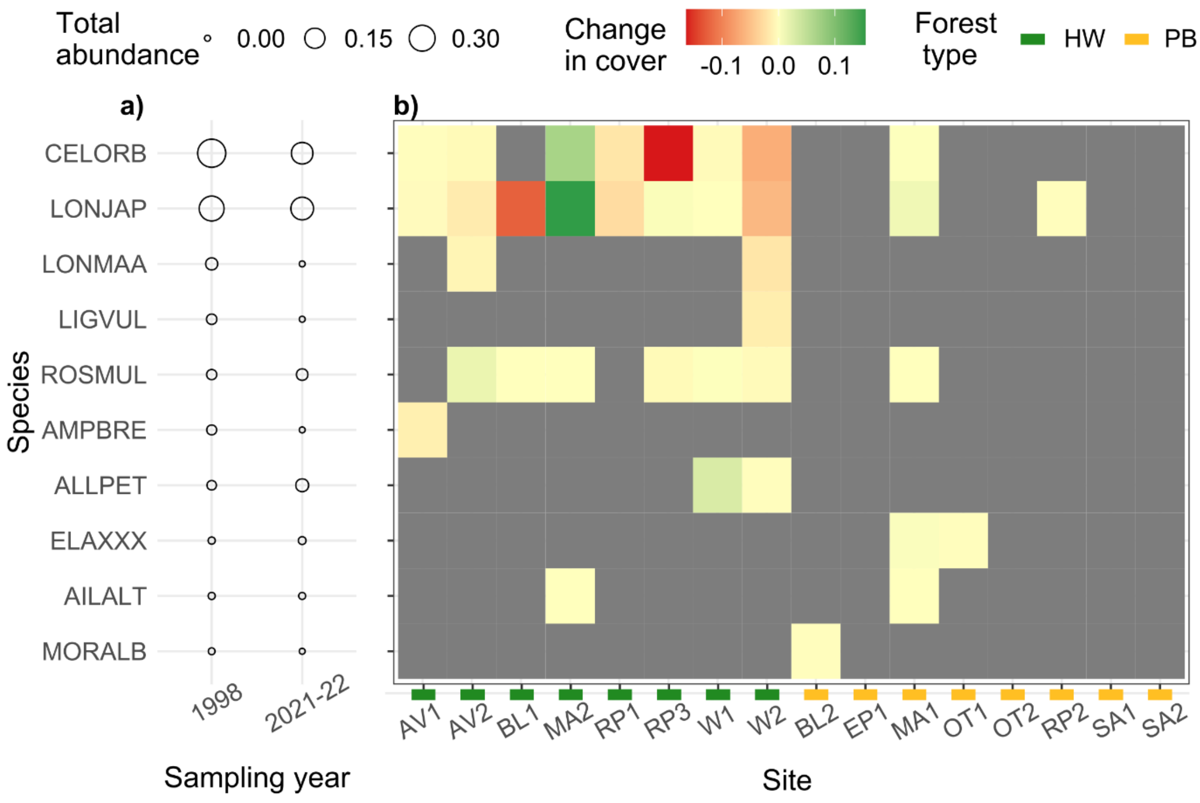

**Figure S6:** Temporal changes in the abundance of the 10 most common introduced species in 1998. a) Total abundance (sum of cover across sites) of each species in 1998 and 2021-22. b) Change in the cover of each species in each site (cover in 1998 – cover in 2021-22). Sites are arranged by forest type. Positive values (light green) indicate increase, and negative values (red) indicate decrease over time. Grey fill indicates absence of the given species in the given site at both time points. From the 1998 dataset, only the data from resampled sites was used. Species names associated with the six-letter codes in a) are shown in table S10.

**Table S1:** Results of mixed-effects ANOVAs testing whether the degree of invasion (richness and total cover of introduced species) in the two forest types significantly changed over time. Site was the random effect, and forest type (hardwood vs pine barrens) and year (1998 vs 2021-22) were the fixed effects.

| **Response: Square-root transformed introduced species cover** | | | |
|---|---|---|---|
| **Predictor** | **Df** | **F** | **P-value** |
| Year | 1 | 0.21 | 0.66 |
| Forest type | 1 | 28.91 | **$9.77 \times 10^{-5}$*** |
| Forest type x year | 1 | 0.29 | 0.60 |
| **Response: Log-transformed introduced species richness** | | | |
| **Predictor** | **Df** | **F** | **P-value** |
| Year | 1 | 0.18 | 0.68 |
| Forest type | 1 | 25.14 | **$1.89 \times 10^{-4}$*** |
| Forest type x year | 1 | 0.89 | 0.36 |

*$p < 0.05$

**Table S2:** Results of t-tests on soil variables, to test whether they differed significantly between hardwood and pine barrens sites. The two-sided unpaired t-test for unequal variances was used.

| **Soil variable (units)** | **Mean of hardwood sites (± s.d)** | **Mean of pine barrens sites (± s.d)** | **t-statistic** | **Df** | **P-value** |
|---|---|---|---|---|---|
| pH | 3.5 ± 0.2 | 3.4 ± 0.3 | 1.7 | 34.9 | 0.10 |
| Calcium (meq/L) | 0.98 ± 0.78 | 0.22 ± 0.10 | 3.9 | 15.4 | **0.001*** |
| Magnesium (meq/L) | 0.98 ± 0.62 | 0.44 ± 0.62 | 2.6 | 32.3 | **0.01*** |
| Cation Exchange Capacity (meq/100g) | 12.8 ± 2.0 | 11.2 ± 3.24 | 1.7 | 33.7 | $0.09^{+}$ |
| Bray Phosphorus (ppm) | 12.4 ± 7.9 | 8.22 ± 5.46 | 1.8 | 25.4 | $0.08^{+}$ |
| Sand (%) | 73.5 ± 6.7 | 75.2 ± 14.8 | -0.5 | 29.3 | 0.64 |
| Silt (%) | 21.2 ± 5.87 | 19.6 ± 12.5 | 0.5 | 29.9 | 0.60 |
| Clay (%) | 5.3 ± 1.1 | 5.2 ± 2.4 | 0.1 | 29.9 | 0.90 |

$^{+}p < 0.1$, $*p < 0.05$

**Table S3:** Results of max tests on dc-CA models, to test for overall model significance. The max test consists of two permutation tests, one where species attributes (traits or species origin) are permuted across species, and the other where site attributes (soil variables) are permuted across sites. The more conservative of the two tests is then used to determine model significance.

| | | | | | | |
|---|---|---|---|---|---|---|
| **1998, model w. functional traits and soil variables as predictors** | | | | | | |
| | **Species-level permutation test** | | | **Site-level permutation test** | | |
| | Df | F | P-value | Df | F | P-value |
| Model | 7 | 4.84 | 0.002* | 6 | 2.25 | 0.01* |
| Residual | 47 | | | 30 | | |
| **1998, model w. species origin as predictor and traits as conditioning variables** | | | | | | |
| | **Species-level permutation test** | | | **Site-level permutation test** | | |
| | Df | F | P-value | Df | F | P-value |
| Model | 1 | 2.19 | 0.37 | 6 | 1.72 | 0.16 |
| Residual | 46 | | | 30 | | |
| **2021-22, model w. functional traits and soil variables as predictors** | | | | | | |
| | **Species-level permutation test** | | | **Site-level permutation test** | | |
| | Df | F | P-value | Df | F | P-value |
| Model | 6 | 6.38 | 0.001* | 6 | 1.84 | 0.03* |
| Residual | 56 | | | 9 | | |
| **2021-22, model w. species origin as predictor and traits as conditioning variables** | | | | | | |
| | **Species-level permutation test** | | | **Site-level permutation test** | | |
| | Df | F | P-value | Df | F | P-value |
| Model | 1 | 1.70 | 0.67 | 6 | 1.01 | 0.47 |

| | | | | |
|---|---|---|---|---|
| Residual | 55 | | | 9 |

*$p < 0.05$

**Table S4:** Results of max tests on individual dc-CA axes from each dc-CA model, to identify significant axes of trait-environment relationships. Only the first three axes from each model are shown.

| **1998** | | | | | | | | |
|---|---|---|---|---|---|---|---|---|
| | **Species-level permutation test** | | | | **Site-level permutation test** | | | |
| | Df | $R^2$ | F | P-value | Df | $R^2$ | F | P-value |
| **Axis 1** | 1 | 0.25 | 20.87 | **0.03*** | 1 | 0.19 | 8.09 | **0.02*** |
| Axis 2 | 1 | 0.08 | 6.26 | 0.36 | 1 | 0.06 | 2.58 | 0.42 |
| Axis 3 | 1 | 0.05 | 4.05 | 0.66 | 1 | 0.04 | 1.72 | 0.64 |
| **2021-22** | | | | | | | | |
| | **Species-level permutation test** | | | | **Site-level permutation test** | | | |
| | Df | $R^2$ | F | P-value | Df | $R^2$ | F | P-value |
| **Axis 1** | 1 | 0.23 | 22.00 | **0.01*** | 1 | 0.32 | 6.33 | **0.01*** |
| Axis 2 | 1 | 0.07 | 6.43 | 0.73 | 1 | 0.09 | 2.06 | 0.70 |
| Axis 3 | 1 | 0.06 | 5.95 | 0.73 | 1 | 0.09 | 2.09 | 0.70 |

*$p < 0.05$

**Table S5:** Results of two-sample Kolmogorov-Smirnov tests on the species pool distributions of continuous traits, to test whether they significantly differed between native and introduced species.

| **Trait** | **KS statistic** | **P-value** |
|---|---|---|
| SLA | 0.2 | 0.38 |
| Height | 0.4 | **0.02*** |
| Leaf C:N ratio | 0.3 | 0.18 |

**Table S6:** Results of t-tests on species pool means of continuous traits, to test whether they significantly differed between native and introduced species.

| **Trait** | **T-statistic** | **Df** | **P-value** |
|---|---|---|---|
| SLA | -1.1 | 29.2 | 0.29 |
| Height | 0.8 | 40.7 | 0.43 |
| Leaf C:N ratio | 2.6 | 40.6 | **0.01*** |

**Table S7:** Results of F-tests on species pool variances of continuous traits, to test whether they significantly differed between native and introduced species pools.

| **Trait** | **F-statistic** | **Df (native, introduced)** | **P-value** |
|---|---|---|---|
| SLA | 1.0 | 48, 16 | 0.87 |
| Height | 1.9 | 50, 17 | 0.16 |
| Leaf C:N ratio | 2.2 | 43, 15 | $0.08^{+}$ |

$^{+}p < 0.1$, $*p < 0.05$

**Table S8:** Results of Chi-squared tests on species pool distributions of categorical traits, to test whether they significantly differed between native and introduced species pools.

| **Trait** | **Chi-squared statistic** | **Df** | **P-value** |
|---|---|---|---|
| Growth Rate | 2.4 | 2 | 0.30 |
| Life form | 2.8 | 3 | 0.42 |
| Shoot C:N ratio | 0.3 | 2 | 0.86 |
| Fire tolerance | 1.4 | 3 | 0.71 |

**Table S9:** Results of permutational MANOVAs testing whether the taxonomic and functional composition of resampled sites significantly changed over time, with separate tests for native and introduced species assemblages. Permutations were restricted to within site.

| **Taxonomic composition** | | | | | | | | |
|---|---|---|---|---|---|---|---|---|
| Predictor | **Native** | | | | **Introduced** | | | |
| | Df | $R^2$ | F | P-value | Df | $R^2$ | F | P-value |
| Sampling year | 1 | 0.06 | 1.75 | **0.001*** | 1 | 0.04 | 1.23 | **0.04*** |
| Residual | 30 | 0.94 | | | 30 | 0.96 | | |
| **Functional composition** | | | | | | | | |
| Predictor | **Native** | | | | **Introduced** | | | |
| | Df | $R^2$ | F | P-value | Df | $R^2$ | F | P-value |
| Sampling year | 1 | 0.03 | 0.89 | 0.15 | 1 | 0.02 | 0.53 | 0.64 |
| Residual | 30 | 0.97 | | | 30 | 0.98 | | |

*$p < 0.05$

**Table S10:** Species names associated with the six-letter codes shown in figures S5-S6.

| Code | Species name | Origin |
|---|---|---|
| ACERUB | *Acer rubrum* | Native |
| AILALT | *Ailanthus altissima* | Introduced |
| ALLPET | *Alliaria petiolata* | Introduced |
| AMPBRE | *Ampelopsis glandulosa* | Introduced |
| BETNIG | *Betula nigra* | Native |
| CELORB | *Celastrus orbiculatus* | Introduced |
| CORFLO | *Cornus florida* | Native |
| ELAXXX | *Elaeagnus angustifolia/Elaeagnus umbellata* | Introduced |
| EURDIV | *Eurybia divaricata* | Native |
| GAUPRO | *Gaultheria procumbens* | Native |
| GAYBAC | *Gaylussacia baccata* | Native |
| JUNVIR | *Juniperus virginiana* | Native |
| LIGVUL | *Ligustrum vulgare* | Introduced |
| LONJAP | *Lonicera japonica* | Introduced |
| LONMAA | *Lonicera maackii* | Introduced |
| MORALB | *Morus alba* | Introduced |

| Code | Species name | Origin |
|---|---|---|
| PARQUI | *Parthenocissus quinquefolia* | Native |
| PINRIG | *Pinus rigida* | Native |
| PRUSER | *Prunus serotina* | Native |
| QUEALB | *Quercus alba* | Native |
| QUEILI | *Quercus ilicifolia* | Native |
| QUEP_C | *Quercus palustris/Quercus coccinea* | Native |
| QUER_V | *Quercus rubra/Quercus velutina* | Native |
| ROSMUL | *Rosa multiflora* | Introduced |
| RUBFLA | *Rubus flagellaris* | Native |
| SASALB | *Sassafras albidum* | Native |
| TOXRAD | *Toxicodendron radicans* | Native |
| VACXXX | *Vaccinium angustifolium/Vaccinium pallidum* | Native |
| VIBACE | *Viburnum acerifolium* | Native |
| VITXXX | *Vitis spp.* | Native |

### Appendix S2: Further details on the obtaining and pre-processing of trait data

*Data sources and pre-processing*

Trait data was gathered from the TRY Plant Trait Database (https://www.try-db.org; Kattge et al., 2020) and the USDA Plants Database (https://plants.usda.gov/; USDA-NRCS, 2023). Data on SLA, leaf C:N ratio, and woodiness (categorical, used for determining life form) was obtained from TRY. Three measures of SLA were available on this database: SLA measured with leaf petiole intact (SLA - petiole included), SLA measured after removing leaf petiole (SLA - petiole excluded), and SLA data where the leaf petiole inclusion/exclusion status was unknown (SLA - petiole unknown). The second of these measures is considered to be the most appropriate measure of SLA (Pérez-Harguindeguy et al., 2013), but more data was available for the other two measures. Therefore, we used SLA - petiole excluded where available. For species with data for one (or both) of the other two measures only, we predicted SLA - petiole excluded through a linear regression of the same against the other two SLA measures.

Data on growth rate, fire tolerance, and life form (referred to as 'growth habit' in the database) came from USDA Plants. Life form categories were reduced to the following four categories: herbs (including forbs, graminoids, and herbaceous vines), lianas (woody vines), shrubs, and trees. Species categorized as 'subshrub' were reclassified into either shrub or herb, based on whether they were woody or not, respectively, using the woodiness trait from TRY. Similarly, vines were placed in either the liana or herb category based on their woodiness. For species labeled as 'shrub/tree' in the database, a height cut-off of 5 m was used to categorize them as either shrub or tree. 'Shrub/vine' species were considered to be lianas.

Height data was obtained from both TRY and USDA Plants. However, the height data in TRY appeared to be inaccurate, and therefore, we used the USDA Plants data where available. For species with height data available on TRY but not on USDA Plants, we carried out a linear regression of the USDA height data against the TRY height data, and predicted the former from the latter. Shoot C:N ratio was also obtained from both databases. For this trait, USDA data was used wherever available, and if not available, TRY data was used.

For each categorical trait, we ensured that roughly equal numbers of species were present in each level of that trait. This was found to be true for all categorical traits except shoot C:N ratio. For that trait, very few species were in the 'low' category, and therefore, the low and medium categories were combined into 'low-medium'.

Traits of species aggregates were calculated by taking the mean (or mode, for categorical traits) of trait values of the constituent species.

*Trait imputation*

For continuous traits, we first carried out stepwise regressions of each trait against all other traits, to select significant predictors. Then, missing values were predicted using linear regressions of the respective trait against the significant predictors. For categorical traits other than life form, missing values were replaced by the most common value in that species' life form. This approach was used because all categorical traits significantly differed between life forms, as determined by Chi-squared tests. All species with missing life form data also did not have data for any other traits, and were removed from the analysis (thus no imputation was necessary for this trait). Imputed data was only used for dc-CA analyses.

*References*

Kattge, J. *et al.* (2020) 'TRY plant trait database – enhanced coverage and open access', *Global Change Biology*, 26(1), pp. 119–188. doi: 10.1111/GCB.14904

Pérez-Harguindeguy, N. A. *et al.* (2013) 'New handbook for standardised measurement of plant functional traits worldwide', *Australian Journal of Botany*, 61, pp. 167–234. doi: 10.1071/BT12225

United States Department of Agriculture National Resources Conservation Service (USDA -NRCS) (2023) *The PLANTS Database*. Available at: https://plants.usda.gov/ (Accessed: 9 March 2023).
